# A greenhouse-based high-throughput phenotyping platform for identification and genetic dissection of resistance to Aphanomyces root rot in field pea

**DOI:** 10.1101/2022.08.01.502415

**Authors:** Md. Abdullah Al Bari, Dimitri Fonseka, John Stenger, Kimberly Zitnick-Anderson, Sikiru Adeniyi Atanda, Hannah Worral, Lisa Piche, Jeonghwa Kim, Mario Morales, Josephine Johnson, Rica Amor Saludares, Paulo Flores, Julie Pasche, Nonoy Bandillo

**Affiliations:** Department of Plant Sciences, North Dakota State University, Fargo, ND 58108-6050, USA; Department of Plant Pathology, North Dakota State University, Fargo, ND 58108-6050, USA; Department of Agricultural and Biosystem Engineering, North Dakota State University, Fargo, ND 58108-6050, USA; North Dakota State University North Central Research Extension Center, 5400 Highway 83 South, Minot, ND 58701, USA

**Keywords:** High-throughput phenotyping, Genome-wide association mapping, Aphanomyces root rot, *Aphanomyces euteiches*, Field pea

## Abstract

Aphanomyces root rot (ARR) is a devastating disease in field pea *(Pisum sativum* L.) that can cause up to 100% crop failure. Assessment of ARR resistance can be a rigorous, costly, time-demanding activity that is relatively low-throughput and prone to human errors. These limits the ability to effectively and efficiently phenotype the disease symptoms arising from ARR infection, which remains a perennial bottleneck to the successful evaluation and incorporation of disease resistance into new cultivars. In this study, we developed a greenhouse-based high throughput phenotyping (HTP) platform that moves along the rails above the greenhouse benches and captures the visual symptoms caused by *Aphanomyces euteiches* in field pea. We pilot tested this platform alongside with conventional visual scoring in five experimental trials under greenhouse conditions, assaying over 12,600 single plants. Precision estimated through broad-sense heritability (*H*^2^) was consistently higher for the HTP-indices (*H*^2^ Exg =0.86) than the traditional visual scores (*H*^2^ DSI=0.59), potentially increasing the power of genetic mapping. We genetically dissected variation for ARR resistance using the HTP-indices, and identified a total of 260 associated single nucleotide polymorphism (SNP) through genome-wide association (GWA) mapping. The number of associated SNP for HTP-indices was consistently higher with some SNP overlapped to the associated SNP identified using the visual scores. We identified numerous small-effect QTLs, with the most significant SNP explaining about 5 to 9% of the phenotypic variance per index, and identified previously mapped genes known to be involved in the biological pathways that trigger immunity against ARR, including Psat5g280480, Psat5g282800, Psat5g282880, and Psat2g167800. We also identified a few novel QTLs with small-effect sizes that may be worthy of validation in the future. The newly identified QTLs and underlying genes, along with genotypes with promising resistance identified in this study, can be useful for improving a long-term, durable resistance to ARR.

## Introduction

Field pea *(Pisum sativum* L., 2n = 14), also known as “dry pea,” is the second most important grain legume in the world, after the common bean *(Phaseolus vulgaris* L.) (FAO, 2016). It is currently grown in most temperate regions of the world for human consumption, livestock feed, and the pet food market. Field pea is an excellent source of proteins, minerals, starches, and dietary fibers (Dahl et al. 2012; McDermott and Wyatt 2017), and its nutritional profile complements cereal grains (Foschia et al.2017). Regular consumption of pea can help fight type 2 diabetes and obesity (Dahl et al. 2012). With the growing popularity of plant-based products such as the Beyond Burger^®^ (https://www.beyondmeat.com) and Ripple’s^®^ non-dairy ice cream and milk options (https://www.ripplefoods.com/ripptein/) (Foschia et al. 2017; Bandillo et al. 2021), which all utilize pea protein as a major constituent, the need for the development of competitive high-protein pea cultivars also increases. In addition to all the benefits promoted by pea consumption, its addition into cropping system can enhance soil health attributes, reduce greenhouse gas emissions, improve soil biota diversity, leading to more sustainable agricultural practices (Magrini et al. 2018). As with any other crop, however, the successful cultivation of field pea can be threatened by abiotic and biotic stressors such as emerging pests and diseases.

Aphanomyces root rot (ARR), caused by *Aphanomyces euteiches* Drechs., is one of the most destructive pea diseases, causing up to 100% crop failure (Gaulin et al. 2007; Gossen et al. 2016; Zitnick-Anderson et al. 2021). In North Dakota, which is the primary target environment of the North Dakota State University (NDSU) pulse breeding program, the disease was first identified in 2014, and since then has been prevalent in pulse-growing regions across North America (Heyman 2008; Sivachandra Kumar et al., 2021; Zitnick-Anderson and Pasche 2016). *Aphanomyces euteiches* has complex genetics and exhibits variable degrees of aggressiveness (Grünwald and Hoheisel 2006; Le May et al. 2018). The symptoms of ARR initially appear as water-soaked and grayish discoloration. As the disease progresses, the roots exhibit honey-brown discoloration and become soft (Gossen et al. 2016; Zitnick-Anderson et al. 2021), lateral and root hairs start to decay, and in advanced stages the entire root mass can turn blackish and decays (Gossen et al. 2016). The symptoms can also extend to the epicotyl giving the lower stem a shrunken appearance, and further disease development can lead to chlorosis, necrosis, stunting, and death of older leaves, that may progress upward along with the shoot (Heyman 2008; Gossen et al. 2016). This is further exacerbated by soil-borne pathogen that can survive in the soil for ten years without a suitable host, making crop rotation less impactful in reducing disease pressure. Due to the lack of improved cultivars and adequate host resistance, chemical control alone falls short in effectively controlling the disease. In addition, excessive use of chemicals is undesirable due to detrimental environmental effects (Zitnick-Anderson and Pasche 2016). Therefore, the development of resistant pea varieties is the eco-friendliest disease management practice (Nelson et al. 2018).

Phenotyping disease symptoms caused by ARR is rigorous, costly, time-consuming, and subject to human errors. As many pulse breeding programs, the NDSU pulse breeding program routinely screens thousands of genotypes and accessions to identify new sources of resistance to ARR. Therefore, it is important to develop a scalable, efficient, and objective-based disease rating platform to ARR.

Image-based high-throughput phenotyping (HTP) is now a readily feasible option for digitally acquiring complex phenotypic data, thanks, in part, to advancements of sensor technologies and improved high-performance computing capabilities (Araus and Cairns 2014; Ma et al. 2020). Digital RGB cameras have been used to capture disease symptoms in a high-throughput manner in an array of crops, including soybean, tomato, and rice (Sugiura et al. 2016; Mim et al. 2019; Reza et al. 2019). In addition, several HTP platforms including greenhouse, ground-based, and aerial systems have been implemented in other crops, providing a better understanding of the genetics underlying important traits such as lodging (Singh et al. 2019), maturity (Volpato et al. 2021), and economically important diseases (DeSalvio et al. 2022). With the rapid development of low-cost sensors and platforms, HTP platforms hold great potential to become an integral part of applied breeding efforts. However, significant developments in processing, methodology, and downstream analysis of image-derived data are still needed to realize its full potential (Singh et al. 2019).

Several previous studies have indicated the complex genetic architecture underlying ARR (Pilet-Nayel et al. 2002), making breeding efforts for developing resistant field pea varieties more difficult. As many as 52 genomic regions conferring quantitative resistance to ARR have been identified, but these discoveries were made evaluating pathogen and host populations from France and to a much lesser degree the Pacific Northwest, USA (Desgroux et al. 2016). Environmental conditions play a role in the success of a cultivar regionally, but resistance is also dependent on the pathogen population. *Aphanomyces euteiches* has been demonstrated to vary in aggressiveness as well as genetically; therefore, it is imperative that both host and pathogen factors are evaluated on a regional basis (Grünwald and Hoheisel et al 2006; LeMay et al. 2018; Quillévéré-Hamard et al. 2018; Zitnick-Anderson et al. 2021). Combining just seven of the genomic regions for partial resistance into a new cultivar has resulted in decreased ARR severity (Lavaud et al. 2016). However, to date, no cultivar with full resistance has been developed (Quillévéré-Hamard et al. 2018).

This study aimed to develop a greenhouse-based high-throughput phenotyping platform to identify sources of resistance to *Aphanomyces euteiches* in a set of 300 advanced pea lines developed by the NDSU pulse breeding program. Current greenhouse screening systems are effective at identifying resistance to ARR; however, these methods require five to six weeks to obtain screening results. The high-throughput system aims to reduce that time to almost half. The collected images from the developed HTP-platform were used to identify the genetic regions and underlying candidate genes associated with ARR resistance using genome-wide association (GW A) mapping. Using the HTP-indices, we showed that the heritable variation was nearly two-fold higher for HTP-derived indices than the ground truth visual scores of the disease. We were also able to identify important genes that may play a key role in breeding for genetic resistance against ARR in the future.

## Materials and Methods

### Plant materials, experimental design and greenhouse growing conditions

A set of 300 F_4:8_ advanced NDSU field pea lines derived from multiple bi-parental crosses were used in this study. Four commercial cultivars were used as checks, including Arcadia, CDC Striker (Warkentin et al. 2004), DS Admiral (Andersen et al. 2002), and PLP 522 with differential responses to ARR disease. This experiment was conducted at the Dalrymple Research Greenhouse Complex located at the NDSU main campus in Fargo, North Dakota. The experiment was laid out following an augmented row-column design with checks arranged in a diagonal fashion. This experimental design was used to correct for the confounding effects due to the orientation and shape of the greenhouse. In this study, each experimental unit (genotype) was represented by seven small pots, where each pot contained a single plant. Small, deep plastic pots (cell diameter of 3.8 cm, depth of 21 cm, and a volume of 164 cubic centimeter) (Stuewe & Songs, Inc., Tangent, OR, USA) with drainage holes were filled with potting soil mixture Promix PGX (Premier Tech Horticulture, Quakertown, PA, USA). The temperature setting was 76 □ during the day and 66.2 □ at night, with 95% relative humidity, with a light/dark cycle of 16 hours of light and eight hours of dark. The experiment was repeated five times.

### Inoculum preparation and disease inoculation

Eight to 10 yellow corn kernels and 75mL of deionized water were placed into a 250mL Erlenmeyer flask and autoclaved for 20 minutes (Carlson 1965; Zitnick-Anderson et al. 2021). Five agar plugs (approximately 2mm diameter) were transferred from a 7-to 10-old *A. euteiches* culture grown on cornmeal agar (CMA) (1L distilled water and 17g CMA [BBL CMA, BD Biosciences, San Jose, CA, United States) to the autoclaved flask containing corn kernels. After incubation in the dark at 20 to 22°C for 7 to 9 days, the medium was decanted from the mycelium. Mycelium was rinsed twice with 100mL of sterilized tap water. After 3 hours, the final rinse water was decanted and 100ml of 10% sterilized soil to extract water (50g of soil, 450ml of tap water) was added prior to incubating for another three hours. The mycelial mat was transferred to a glass Petri dish (9mm diameter), and 100mL of pea root exudate solution (sterilized tap water) was added (Shang et al. 2000). Pea root exudate solution was prepared by placing 10 pea seeds surface sterilized in a 1% sodium hypochlorite (NaOCl) solution in a sterile 250ml Erlenmeyer flask containing 5ml of sterile distilled water. Flasks were covered with aluminum foil and incubated in dark at room temperature for 14 days. The solution was diluted 1:1 using sterilized tap water prior to use. Cultures in exudate solution were aerated in Petri dishes for eight to 12 hours at room temperature. Zoospore production was routinely monitored every six hours. The solution was filtered through a single layer of cheesecloth and the zoospore concentration was adjusted to 10^4^ zoospores mL^-1^ using a hemocytometer. Field pea plants were inoculated in the greenhouse as soon as zoospore concentration was calculated to avoid encystment of zoospores. Eight to nine days after planting, emerged plants were watered thoroughly and inoculated by pipetting 1mL of the zoospore inoculum to the base of the plants (**Supplementary Table 1**). Plants were not watered until the day following inoculation as to not rinse the inoculum from the soil.

### Development of a high-throughput phenotyping platform and generation of HTP-disease indices

The phenotyping for ARR in dry pea was carried out using images captured from a digital single-lens reflex camera (EOS Rebel T7i, Canon Inc, Tokyo, Japan) (Figure 1). The camera was affixed to a custom cart built to interface and utilize the c-channel in the rafters of the existing greenhouse structure as rails. This allowed the cart to freely transverse the length of the greenhouse room to capture images above each positioned bench containing the sample plants. As the cart moved from bench to bench it was triggered using an iPad (A1584, Apple Inc., Cupertino, CA) and Canon’s Camera Connect application. As there were two c-channels in the room, two carts with cameras were utilized to evaluate the majority of greenhouse space available. Image capture began a week after inoculation and continued for four to eight periods during disease development. The three bands, red (r), green (g), and blue (b), were used to compute vegetation indices for further analysis. Prior to calculating indices, each band was converted from unsigned 8-bit integer values to floating point values then scaled to a unit length and normalized across bands using the following equation (Bannari et al. 1995; Woebbecke et al. 1995):

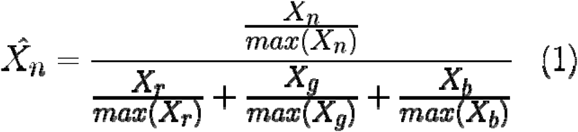

**Figure 1.**
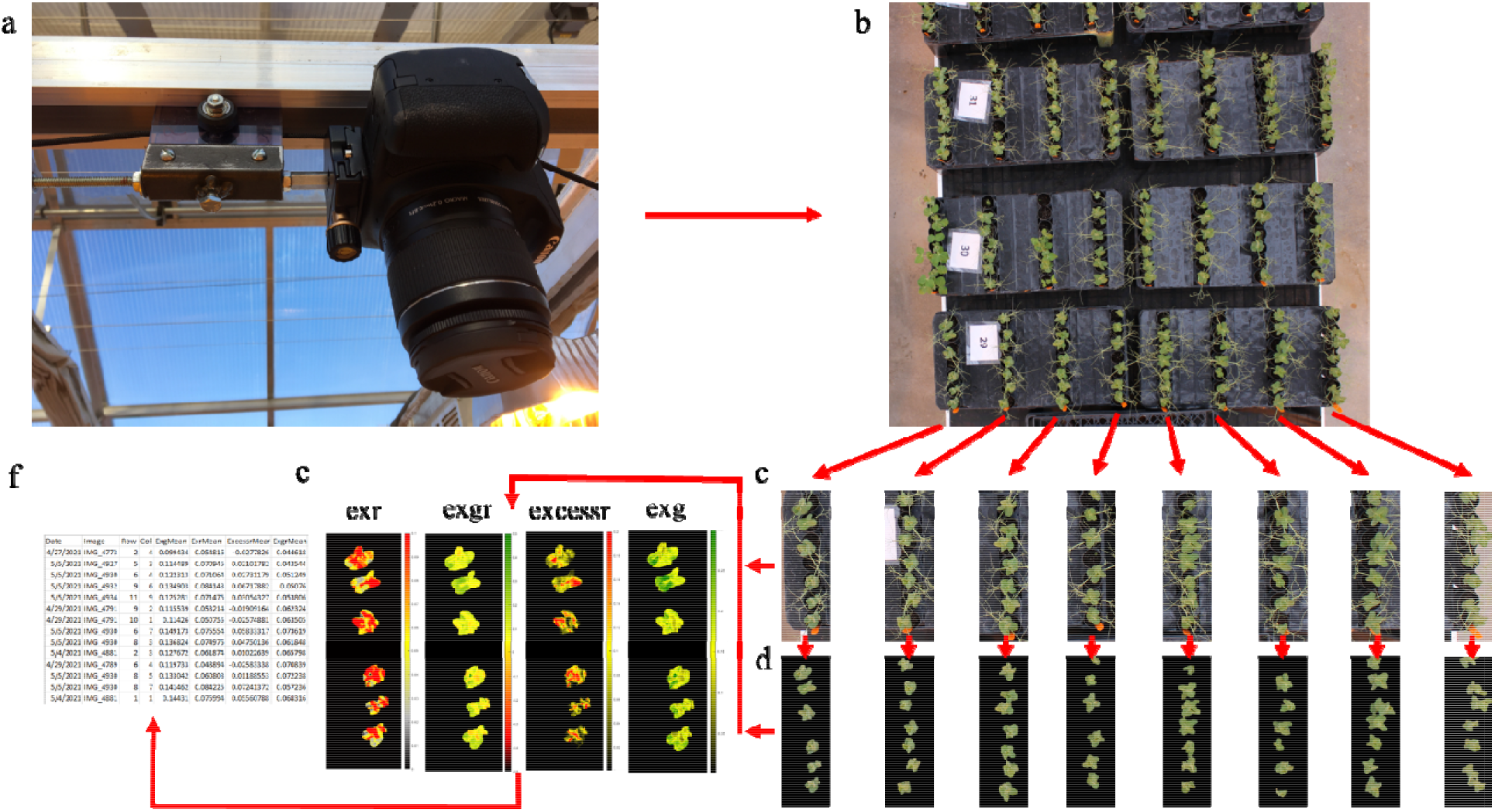
Imagery data collection process overview a) image capture from a digital single-lens reflex camera on custom greenhouse rail mount, b) image captured of multiple experimental units on a bench, c) image rotation and grid extraction of uniform individual experimental units, d) image segmentation through Otsu thresholding and mathematical morphology operations to isolate plant leaves from background and tendrils, e) generation of index values and index masking to obtain mean index values, f) tabulation of results for downstream analysis in a comma-separated value file format.

Where X_n_ is a matrix relating the n^th^ band of the image. X_r_, X_g_, and X_b n_ is a matrix of normalized band values for the n^th^ band of the image utilized for further index computations.

Using the resulting dataset, multiple indices were estimated to describe disease progress differences among tested accessions. Excess green (exg) is a vegetation index which measures the degree to which green dominates over all other colors in the image and was calculated as:

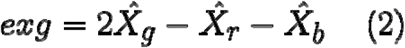

As symptomatic plants are expected to show chlorosis and necrosis, thus lessening the relative ratio of green when compared with other colors, those plants with lower exg values are anticipated to be associated with increased disease symptomology. However, increases in the relative ratio of blue hue are as impactful as increases in red on index values. Plants having higher amounts of blue color, by virtue of their genetics or more intense glabrous cuticular coating, may also have reductions in exg value that would confound the anticipated effects of disease and reduce its usefulness in identification of symptomatic plants.

To reduce the impact of such confounded results, a similar approach was tested using a similar index emphasizing red features. In literature, the term exg is used freely relating to several distinct indices. To separate these confounded terms this index will be referred to as excessr and was calculated as the following (Dutta et al. 2022):

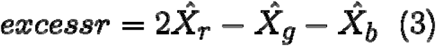

It is anticipated that plants having increased yellow chlorotic or brown necrotic symptomology will have an increased ratio of red to green or blue in their coloration, thus plants having higher excessr values will be considered indicative of plants having increased indications of disease presence and/or severity.

An additional index investigated to evaluate the level of yellowing of plants was the difference in exg and excess red (exr) (Mayer et al. 1998). Exr is a weighted difference of the amount of red to that of green coloration and is calculated as:

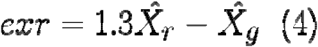

This again, could be a good indication as increases in the relative dominance of red is expected to be associated with plant yellowing and in-turn disease presence and/or severity. The difference in exg to exr (exgr) may also be informative in distinguishing vegetation (Mayer and Neto 2008). To attempt to separate the effects of blue hue from those of red and the index exgr was calculated as:

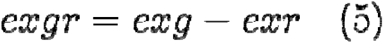

Mayer and Neto (2008) found the index to be more useful in defining green vegetation from background elements containing yellow and dead straw tissue, which is a similar problem that we were facing on this study. Those plants having relatively high ratios of green to red should obtain high index values, while those demonstrating yellow chlorotic or brown necrotic symptomology should result in relatively low values. Less emphasis is placed on the blue coloration as red has a comparatively higher weight. It is anticipated that this would provide more accurate results as compared to the exg index, where red and blue are equally weighted. The maximum respective indices of different time points were used for the analyses.

### Image processing workflow

Entire experimental benches each having trays of rows of plants were captured in a single image. Each row of plant pots relating to an accession was considered an experimental unit and was extracted and evaluated jointly for each collection date. A custom MATLAB (R2020b, MathWorks, Natick MA) program was created to rotate tine selected image to vertically align rows and extract experimental units from their grid arrangement on the bench and label the extracted experimental units with trial identification numbers.

Pixel data extracted from each experimental unit was then processed by custom Python script to extract average index values for each experimental unit on each date, and to output a csv file containing the tabular data. For each experimental unit, a masked image was created as a combination of exg and excessr indices. Otsu thresholding (Otsu 1979) was used to extract the plants from the background. An automated process utilizing mathematical morphological operations was used to remove fine features, such as tendrils, such that the majority of the data was extracted from leaf tissue (Haralick et al. 1987). For each dataset, a small number of experimental units failed to segment correctly utilizing this automated procedure. For these experimental units, generated masks were manually adjusted prior to data extraction. The binary mask was applied to all calculated index images for that experimental unit, such that mean values obtained resulted from sample plant foliage within the image. Generated mean values for each experimental unit for each investigated index were used for subsequent evaluation of genetic effects.

### Visual disease scoring

Conventional scoring was conducted to compare with scores derived from the HTP setup. Plants were harvested within 28 to 39 days after inoculation (**Supplementary Table 1**). Plants were gently lifted from the pots and potting medium and carefully removed without damaging the roots. The roots were then washed with slightly warm tap water (**Figure 2**). Disease scores for pea ARR severity were measured on a scale of 0 to 5; where 0 = healthy plant, 1 = slight discoloration, 2 = moderate discoloration with some shriveling root hairs, 3 = extensive discoloration but epicotyl is not shrunken, 4 = extensive discoloration with a shrunken epicotyl, and 5 = root is partially or completely rotted and detached from the plant. Using the scores from seven plants, we generated two visually derived scores: 1) disease severity index (DSI) and 2) maximum score (Max_score). The DSI was generated following the equation:

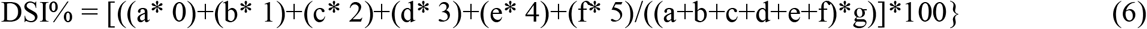

where a, b, c, d, e and f represent the number of plants with the disease severity ratings of 0, 1, 2, 3, 4 and 5 respectively, and g represents the highest root rot severity rating (McKinney 1923; Li et al. 2014). The Max_score was generated to capture the potential maximum disease severity and was extracted using the highest score value among the seven plants evaluated on each experimental unit.

**Figure 2.**
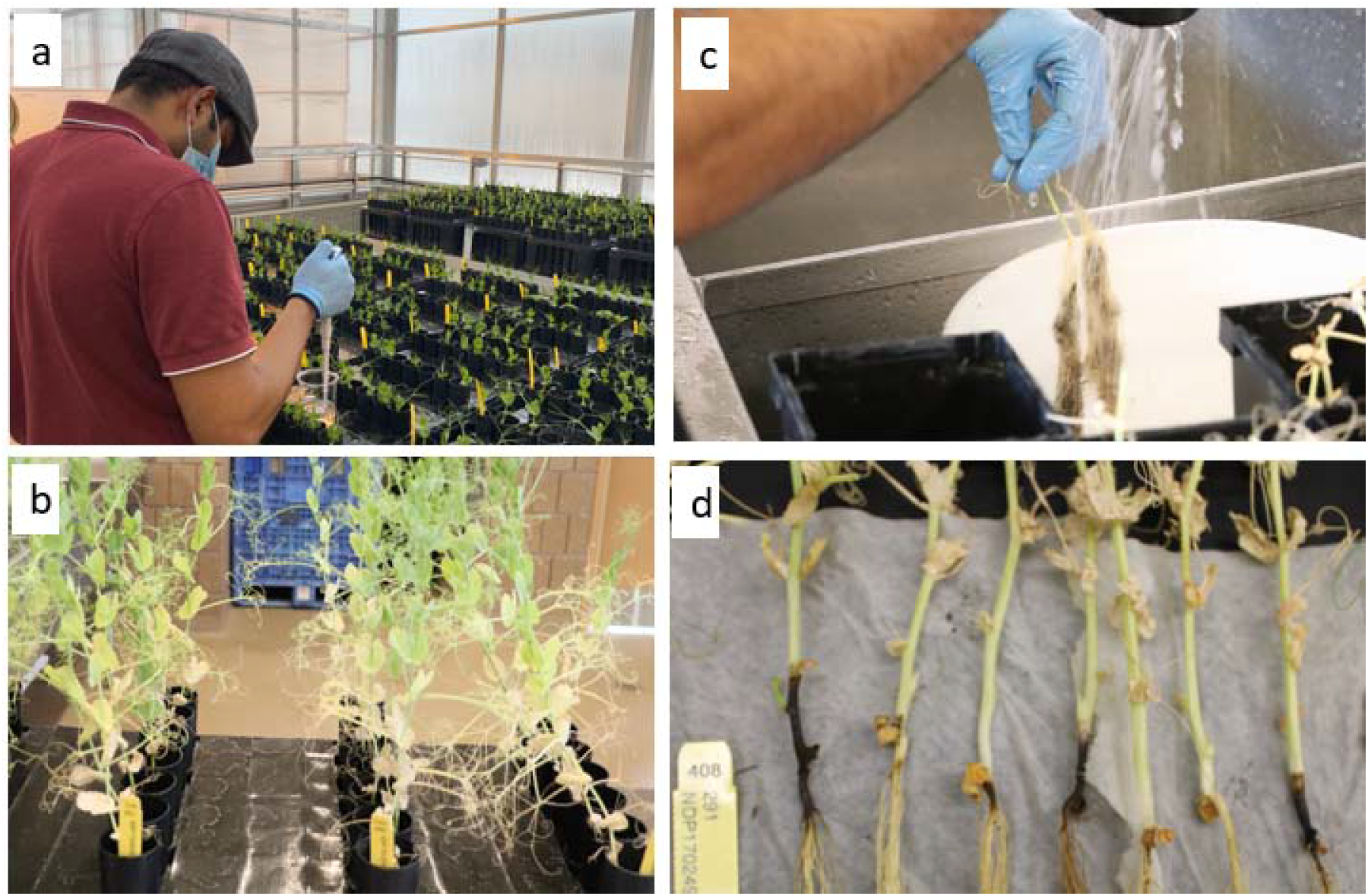
The step-by-step procedures for visual scoring of ARR resistanc: a) inoculation of 7 to 9-day old seedlings, b) harvesting, c) rinsing of roots, d) visual scoring *Aphanomyces* root rot.

### Statistical analyses of visual score and HTP-indices

We fit a mixed model to analyze the data using a two-stage approach in ASReml-R package (Butler et al. 2017). In the first stage, each trial (e.g., replicate) was independently analyzed to estimate the best linear unbiased estimates (BLUE) of each line. The model was fitted following the linear mixed model below.

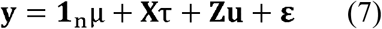

where **y** (n×1) is the vector of visual score or HTP-indices of the lines, μ is the overall mean, **X** and **Z** are the design matrices for fixed and random effects, *τ* is the fixed effect of genotypes, u is the random effect of the row and ε is the residual error. The variance of the random term and the residual variance can be expressed as follows:

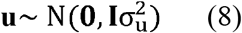

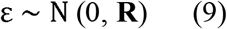

where **I** is an identity matrix, 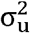 is the variance of the row effect and **R** is the separable autoregressive (AR) variance-covariance matrix defined as:

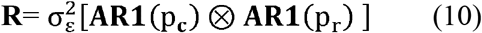

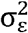 is the spatial residual variance and **AR1**(p_c_) ⊗ **AR1**(p_r_) is the Kronecker product of first-order autoregressive processed across columns and rows, respectively.

The adjusted BLUEs and respective weights from stage one analysis of individual trials were retrieved and then used as inputs for the second stage across replicates of trial over time to obtain BLUPs.

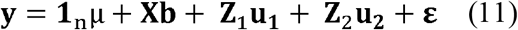

where **y** (n×1) is the adjusted BLUEs of the genotypes implied by the weights in the trials (1…5), μ is the overall mean, b is the fixed effect of the trial replicates. **u_1_** is the random effect of the genotypes with 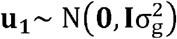, **I** is the identity matrix and 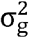 is the genetic variance. **u_2_** is the random effect of genotype by trial interaction, with 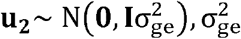 is the variance of the random effect. **ε** is the residual error, with 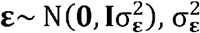 is the residual variance. **X** and **Z** are the design matrix for fixed and random effects as mentioned above.

The broad-sense heritability of each trait was calculated using the equation (12) with variance components derived from equation (7):

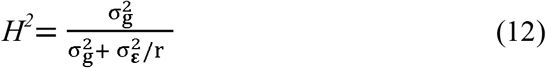

where 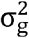 is genetic variance, 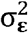 is error variance, and r is number of replication.

### DNA extraction, genotyping-by-sequencing and calling single nucleotide polymorphisms

Young, actively growing leaf tissue was harvested from plants, growing within the greenhouse. Total DNA was extracted from fresh and/or lyophilized leaf tissue using the DNeasy^®^ Plant Mini Kit (Qiagen, Germantown, MD) according to the manufacturer’s instructions and eluted in 100μl. The resultant DNA samples were quantified using the Qubit dsDNA BR Assay kit and Qubit 4.0 fluorometer (Life Technologies Corporation, Eugene, OR).

All DNA samples were standardized to a final concentration of 25 ng/μl and sent to a genomic center for genotyping-by-sequencing (GBS). Dual-indexed GBS libraries were prepared using restriction enzyme ApeKI (Elshire et al. 2011), and combined into a single pool and sequenced across 1.5 lanes of a NovaSeq S1×100-pb run. This generated ≈ 1,000 M pass filter reads for the pool. The mean quality scores ≥ Q30 were obtained for all libraries. The resulting quality reads were aligned to the pea reference genome (Kreplak et al. 2019).

The Single Nucleotide Polymorphism (SNP) marker dataset consisted of 28,832 markers, any unanchored markers were removed leaving a SNP data set of 25,816 markers. From the SNP marker dataset, minor allele frequency values (MAF < 0.01) less than 1% were removed, heterozygosity values of more than 80% were removed, imputation was carried out using the *k-* nearest neighbor genotype imputation method (Money et al. 2015) from the marker matrix. The final filtered genotypic data set consisted of 11,833 curated markers that were used for downstream analyses.

### Genome-wide association mapping

Using visual score and HTP-disease indices as phenotypes, GWA mapping was performed in a set of ~300 advanced NDSU pea lines genotyped with filtered SNP dataset comprised of 11,833 SNP markers. To control spurious association, population structure was accounted for by spectral decomposition of the genomic relationship matrix of the lines. The first three principal components were used in the standard GWA mixed linear model (MLM):

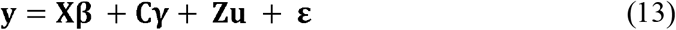

where **y** is a vector of the BLUPs, **X, C** and **Z** are design matrix for **β, γ** and **u**, respectively. **β** is a vector of fixed marker effects; ***γ*** is a vector of subpopulation effects; ***u*** is a vector of the random additive genetic background effects associated with the lines. In addition, general linear model (GLM), compressed mixed linear model (CMLM), and fixed and random model circulating probability unification (FarmCPU) were also implemented.

A comparison-wise error rate was calculated by the Li and Ji (2005) method to control the experiment-wise error rate. The effective number of independent tests (Meff) were calculated from the correlation matrix and eigenvalue decomposition of ~12 K SNP markers. The test criteria were adjusted with the following equation (Šidák 1967):

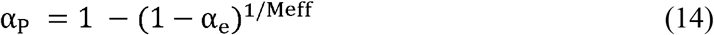

where, α_P_ is the computed comparison-wise error rate, and α_e_ is the trial-wise error rate. In this study, trial-wise error rate is α_e_= 0.05. Experiment wise error rate is used to control type I error due to multiple testing by adjusting alpha value α_e_ = 0.05 to α_P_ = (0.05/473) = 0. 00011, where 473 was the effective number of tests. Several SNPs were in Linkage Disequilibrium (LD) due to physical distance (e.g. linkage), meaning some SNPs were not really independent. Therefore, an effective number of tests (Meff) was used to calculate the p-value.

GWA tables with Li and Ji (2005) threshold at 0.05 experiment-wise error rate of probability i.e., -log_10_ (α_P_) = 3.9 was considered as significant to restrict false marker-trait association (MTA). The GWA results were extracted from GAPIT to further construct Manhattan plots using the CMplot package in R (Yin et al. 2021).

### Candidate gene identification

Candidate genes adjacent to significant SNPs to have insights into the genetic architecture of ARR resistance. The physical region within a ±50 kb window centered on the peak SNPs was chosen to select candidate genes. The effect of SNPs on gene functions within the window was obtained from the annotated file Pisum_sativum_v1a_genes.gff3 (Kreplak et al. 2019).

## RESULTS

### HTP-derived indices have higher heritable variation than visual scores

*H^2^* was consistently higher for HTP-derived indices than visual scores (**Figure 3**). Among HTP indices, exg and exgr had the highest *H^2^* at 0.86 and 0.82, respectively. *H^2^* for visual scores was higher for DSI (*H^2^* = 0.59) than Max_score (*H^2^* = 0.29). Across replications, heritable variation for HTP-derived indices showed nearly two-fold improvement over visual scores. We note that *H^2^* increased with increasing number of replications, indicating the importance of replication when visually scoring ARR which is often subjective and prone to human error.

**Figure 3.**
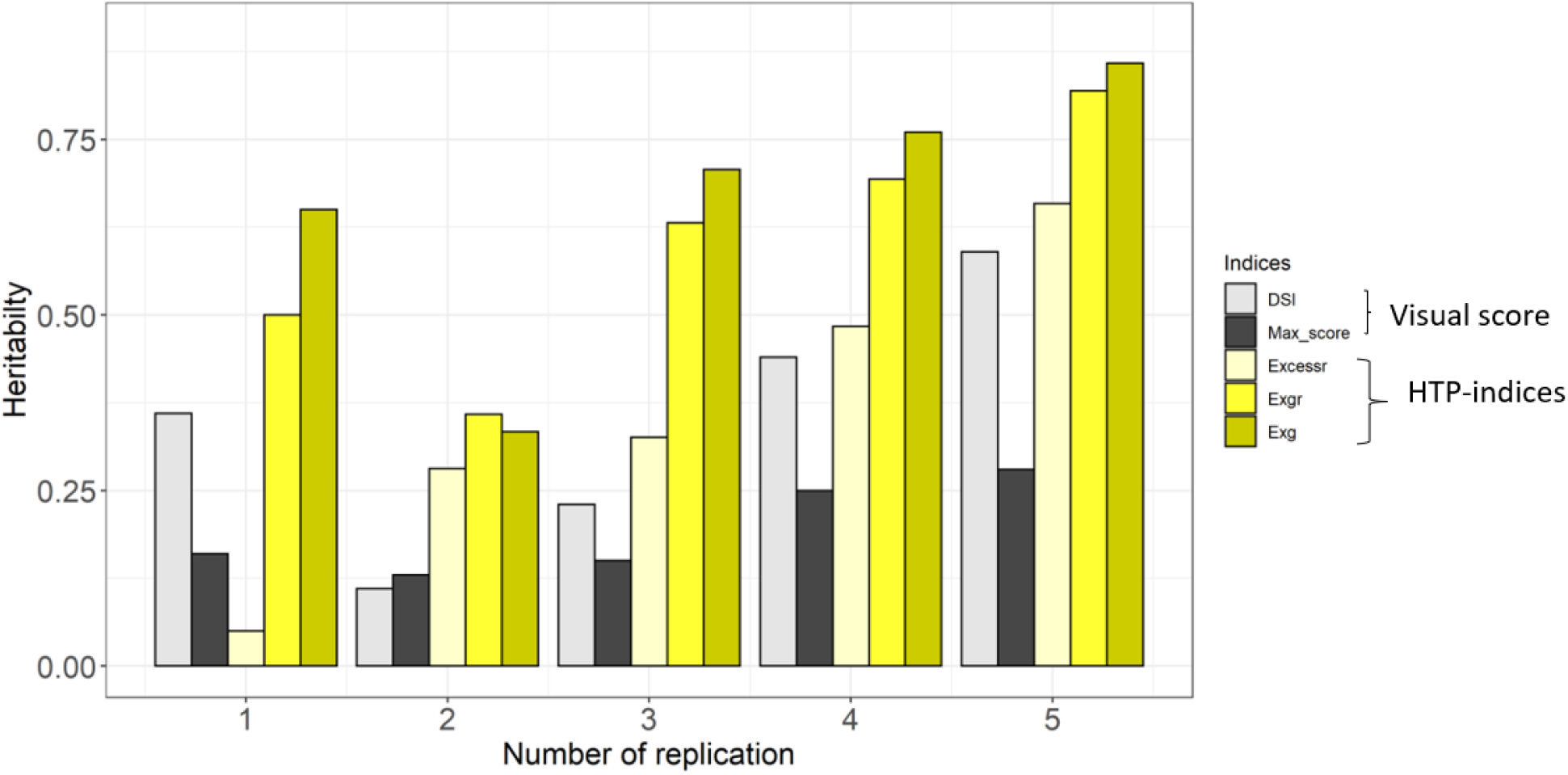
Heritability estimates of visual score (DSI, Max_score), and HTP-indices (Excessr, Exgr, Exg) with increasing number of replications (1 to 5); DSI is the percent disease severity index, Max_score is the maximum recorded score across plants for each line, Excessr is the excess red, Exgr is derived by subtractingexr from exg, and Exg is the excess green.

### Moderate to high correlations among HTP-indices and visual scores

High significant correlations were observed among HTP-indices that ranged from 0.17 to 0.88. Exg and exgr had the highest correlations at 0.88, whereas exg and excessr had modest correlation at 0.17. As expected, we observed high correlations between visual indices. For example, DSI and Max_score had a greater correlation at 0.80. We observed a modest correlation at 0.18 between DSI and HTP-indices (**Supplementary Figures 1 and 2**), which is expected because some genotypes had root rot but showing no visible symptoms on top of the canopy.

### HTP-indices can differentiate genotypes with promising resistance to ARR

We observed a substantial variability on the disease response of 300 advanced breeding lines. Based on DSI, the distribution within and across the five replications ranged from 5 to nearly 80% severity, indicating the severity of disease. We found 23% of genotypes (68 pea lines) outperformed the better moderately resistant reference check Arcadia (DSI = 16%). The results from the combined analyses across replications showed that 47% (140 pea lines) of the advanced breeding lines displayed lower severity compared to the second reference check (partial resistant) DS Admiral (DSI = 20%). About 57% (172 pea lines) of the genotypes outperformed the moderately resistant reference check CDC Striker (DSI = 22%) (**Figure 4, Supplementary Table 3**).

**Figure 4.**
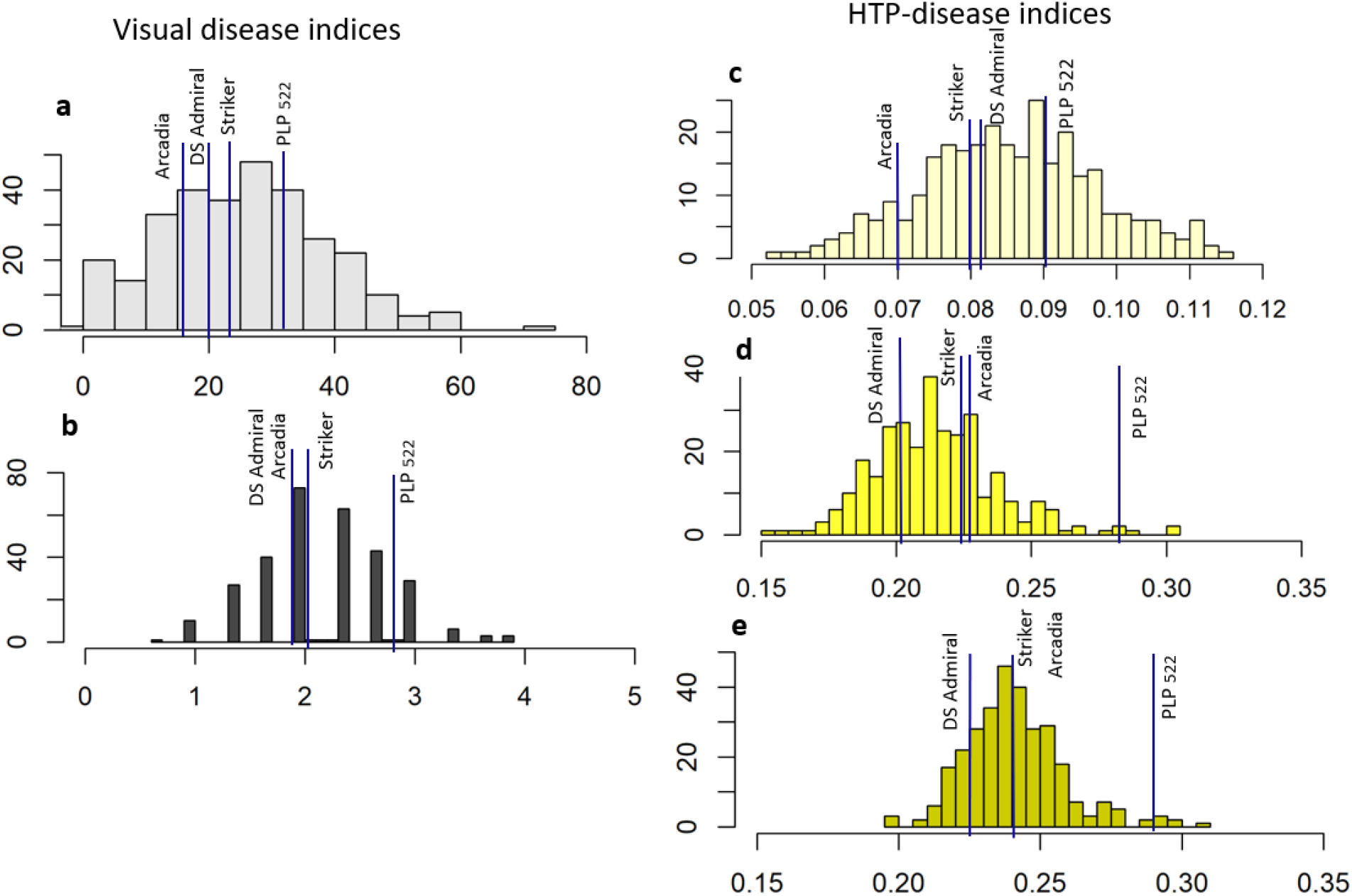
Distribution of visual score and HTP-indices for inference of ARR resistance; The vertical solid blue line were indicated by checks namely: Arcadia, CDC Striker, DS Admiral, and Check 522; a) percent disease severity index (DSI), b) maximum score (Max_score), c) excess red (Excessr), d) Exgr, e) excess green (Exg); y-axis represent frequency.

We explored the utility of HTP-indices for selection of genotypes with promising resistance. The higher exg index values were considered moderately resistant. The best reference check for exg was PLP 522, with an index value of 0.3. For DSI, however, this check had high disease severity (31%) compared to Arcadia (16%), and only two genotypes outperformed the check when using the exg values. The relative performance of the checks (Arcadia, DS Admiral and Striker) was 0.2 for exg, and 43% (128 pea lines) of the genotypes showed higher exg index compared to those checks. Similarly, based on exgr and excessr values, around 34% (102 pea lines) and 41% (122 pea lines), respectively, of field pea genotypes were better when compared to the reference checks (**Supplementary Table 3**). Using both HTP-indices and visual scores, we identified nearly 10% (29/300) of overlapping genotypes that could have promising resistance to Aphanomyces root rot, (**Supplementary Table 4**). The identified genotypes are worthy of further validation in the future.

### GWA mapping provides insight into the genetic architecture of ARR

We genetically dissected variation for ARR resistance using the HTP-indices using the four GWA methods (GLM, MLM, CMLM, FarmCPU), and identified a total of 260 associated SNP using both visual scores and HTP-indices. The number of associated SNP for HTP-indices was consistently higher with some SNP overlapped to the associated SNP identified using the visual scores. A total of 43 significant SNPs was associated for visual scores, including both Max_score and DSI, whereas there were 217 significant SNP associated for HTP-indices (**Supplementary Figure 3**). Most of the signification associated SNP for HTP-indices were located on chromosomes 1, 2, 3, 5, and 7, respectively (**Figure 5, Table 1**). We identified numerous small-effect QTLs, with the most significant SNP explaining about 5 to 9% of the phenotypic variance per index (**Figure 5, Table 1**). We found a substantial overlap (83%) in associated SNP for HTP-disease indices (e.g., exg and exgr), which was likely a result of these variables being strongly correlated with one another, although pleiotropy could play a role. However, only a few associated SNPs overlapped between visual score and HTP-indices, and a few nearby associated SNP potentially in linkage disequilibrium among associated SNPs.

**Figure 5.**
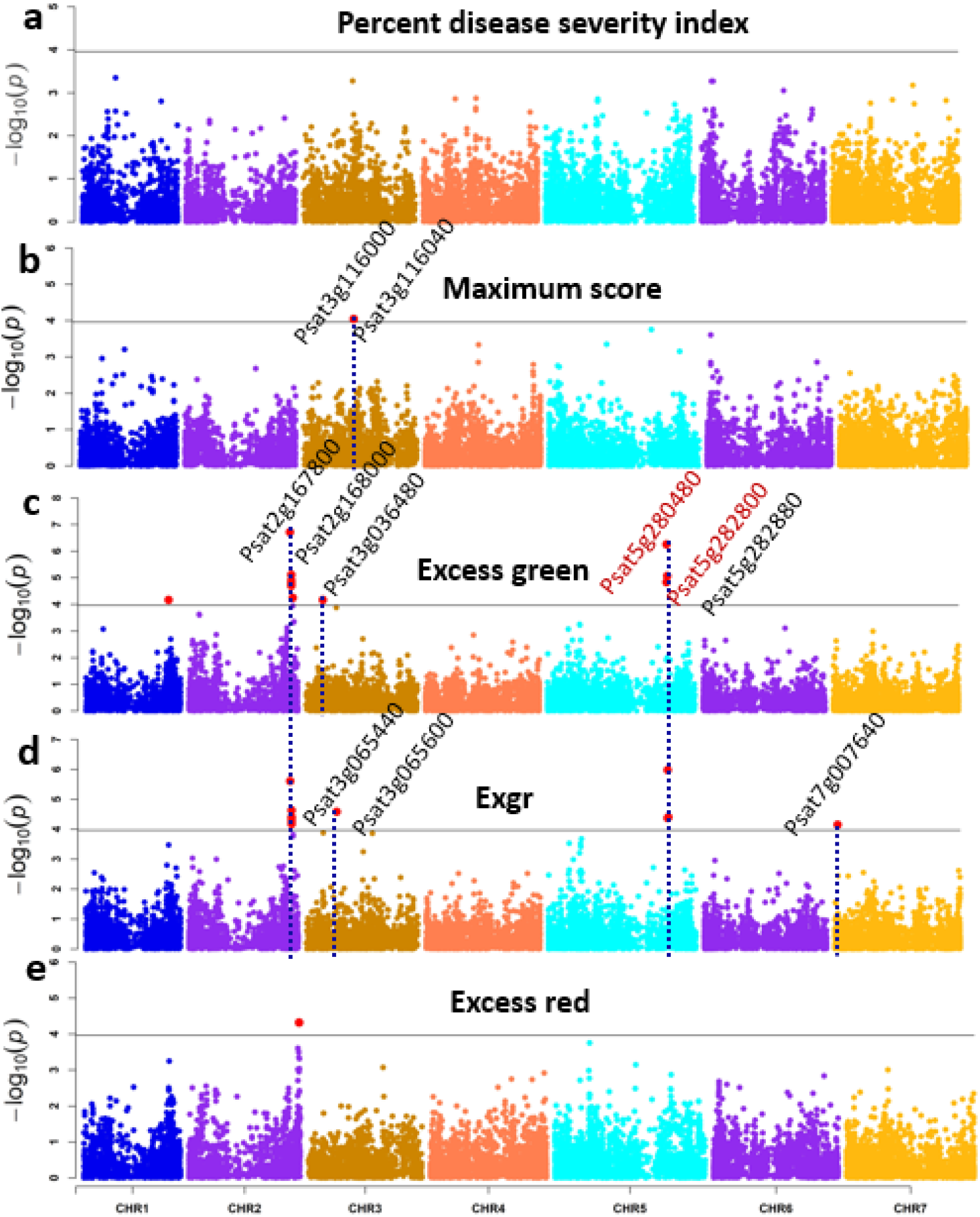
Manhattan plots showing significant SNPs from GWA mapping for a) percent disease severity index (DSI), b) maximum score (Max_score) b) excess green, c) exgr, and d) excess red. Annotated genes known to be associated with ARR resistance near peak SNP region on chromosome 2, 3 and 5 for visual score (Maximum score) and HTP-indices (excess green and exgr). Putatively candidate genes are highlighted in red.

**Table 1.**
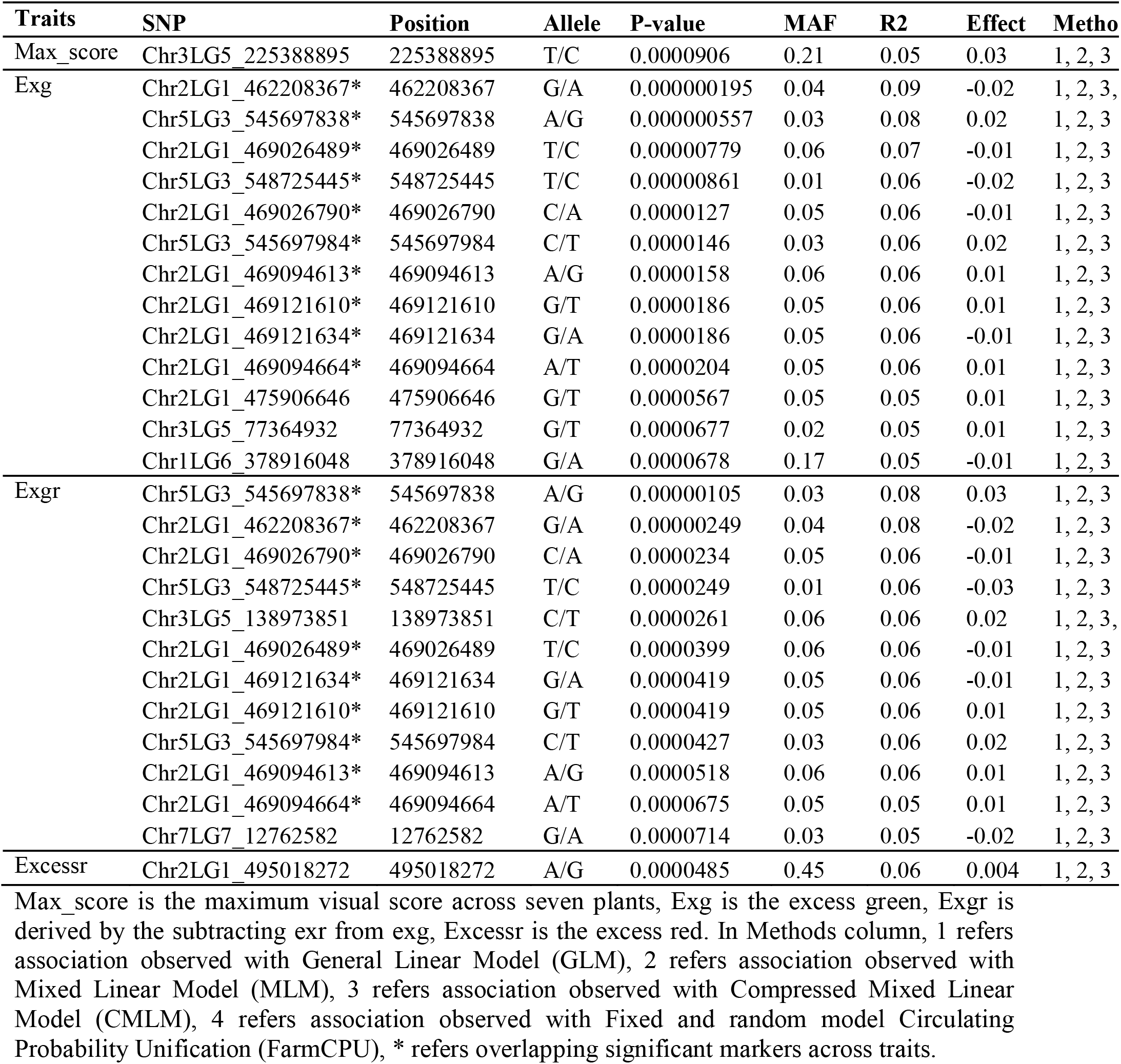
Significant SNPs associated with visual score and HTP-indices.

| Traits | SNP | Position | Allele | P-value | MAF | R2 | Effect | Metho |
| --- | --- | --- | --- | --- | --- | --- | --- | --- |
| Max_score | Chr3LG5_225388895 | 225388895 | T/C | 0.0000906 | 0.21 | 0.05 | 0.03 | 1, 2, 3 |
| Exg | Chr2LG1_462208367* | 462208367 | G/A | 0.000000195 | 0.04 | 0.09 | -0.02 | 1, 2, 3, |
|  | Chr5LG3_545697838* | 545697838 | A/G | 0.000000557 | 0.03 | 0.08 | 0.02 | 1, 2, 3 |
|  | Chr2LG1_469026489* | 469026489 | T/C | 0.00000779 | 0.06 | 0.07 | -0.01 | 1, 2, 3 |
|  | Chr5LG3_548725445* | 548725445 | T/C | 0.00000861 | 0.01 | 0.06 | -0.02 | 1, 2, 3 |
|  | Chr2LG1_469026790* | 469026790 | C/A | 0.0000127 | 0.05 | 0.06 | -0.01 | 1, 2, 3 |
|  | Chr5LG3_545697984* | 545697984 | C/T | 0.0000146 | 0.03 | 0.06 | 0.02 | 1, 2, 3 |
|  | Chr2LG1_469094613* | 469094613 | A/G | 0.0000158 | 0.06 | 0.06 | 0.01 | 1, 2, 3 |
|  | Chr2LG1_469121610* | 469121610 | G/T | 0.0000186 | 0.05 | 0.06 | 0.01 | 1, 2, 3 |
|  | Chr2LG1_469121634* | 469121634 | G/A | 0.0000186 | 0.05 | 0.06 | -0.01 | 1, 2, 3 |
|  | Chr2LG1_469094664* | 469094664 | A/T | 0.0000204 | 0.05 | 0.06 | 0.01 | 1, 2, 3 |
|  | Chr2LG1_475906646 | 475906646 | G/T | 0.0000567 | 0.05 | 0.05 | 0.01 | 1, 2, 3 |
|  | Chr3LG5_77364932 | 77364932 | G/T | 0.0000677 | 0.02 | 0.05 | 0.01 | 1, 2, 3 |
|  | Chr1LG6_378916048 | 378916048 | G/A | 0.0000678 | 0.17 | 0.05 | -0.01 | 1, 2, 3 |
| Exgr | Chr5LG3_545697838* | 545697838 | A/G | 0.00000105 | 0.03 | 0.08 | 0.03 | 1, 2, 3 |
|  | Chr2LG1_462208367* | 462208367 | G/A | 0.00000249 | 0.04 | 0.08 | -0.02 | 1, 2, 3 |
|  | Chr2LG1_469026790* | 469026790 | C/A | 0.0000234 | 0.05 | 0.06 | -0.01 | 1, 2, 3 |
|  | Chr5LG3_548725445* | 548725445 | T/C | 0.0000249 | 0.01 | 0.06 | -0.03 | 1, 2, 3 |
|  | Chr3LG5_138973851 | 138973851 | C/T | 0.0000261 | 0.06 | 0.06 | 0.02 | 1, 2, 3, |
|  | Chr2LG1_469026489* | 469026489 | T/C | 0.0000399 | 0.06 | 0.06 | -0.01 | 1, 2, 3 |
|  | Chr2LG1_469121634* | 469121634 | G/A | 0.0000419 | 0.05 | 0.06 | -0.01 | 1, 2, 3 |
|  | Chr2LG1_469121610* | 469121610 | G/T | 0.0000419 | 0.05 | 0.06 | 0.01 | 1, 2, 3 |
|  | Chr5LG3_545697984* | 545697984 | C/T | 0.0000427 | 0.03 | 0.06 | 0.02 | 1, 2, 3 |
|  | Chr2LG1_469094613* | 469094613 | A/G | 0.0000518 | 0.06 | 0.06 | 0.01 | 1, 2, 3 |
|  | Chr2LG1_469094664* | 469094664 | A/T | 0.0000675 | 0.05 | 0.05 | 0.01 | 1, 2, 3 |
|  | Chr7LG7_12762582 | 12762582 | G/A | 0.0000714 | 0.03 | 0.05 | -0.02 | 1, 2, 3 |
| Excessr | Chr2LG1_495018272 | 495018272 | A/G | 0.0000485 | 0.45 | 0.06 | 0.004 | 1, 2, 3 |

Max\_score is the maximum visual score across seven plants, Exg is the excess green, Exgr is derived by the subtracting exr from exg, Excessr is the excess red. In Methods column, 1 refers association observed with General Linear Model (GLM), 2 refers association observed with Mixed Linear Model (MLM), 3 refers association observed with Compressed Mixed Linear Model (CMLM), 4 refers association observed with Fixed and random model Circulating Probability Unification (FarmCPU), \* refers overlapping significant markers across traits.

### GWA mapping identified candidate genes related to disease resistance

Unless otherwise indicated, we only used the results from the Q+K model for narrowing down the candidate genes. Compared to the other GWA mapping models, the Q□+□K model was chosen for reporting of associated loci because it sufficiently reduced false positives. A search was carried out for annotated genes co-located within ±50 kb of the significant SNPs associated for all traits. Overall, we identified a total of 28 putative genes with annotation generally related to response to biotic stresses, and response to biotic stimulus. Only two genes were narrowed down for the visual scores, with the most significant SNP for the visual score (e.g., Max_score) on chromosome 3 co-localized with Psat3g116000, which known to be involved in plant recognition of pathogens and plants defense mechanisms (**Supplementary Table 5**) (Palma et al. 2007). The gene is also reported to be involved in the regulation of cell wall pectin metabolism, in cellular processes, carbohydrate metabolic pathways, cellular component organization, and other metabolic processes (Geng et al. 2017). Psat3g116040, exhibits a response to abscisic acid, that is reported to be responsive to endogenous stimulus and to other chemicals (Lan et al. 2010).

Twenty-six out of out of 28 candidate genes were the underlying genes identified for the HTP-indices, eight of which were identified in at least two HTP-indices (**Supplementary Table 5**). The overlapping putative genes (**Supplementary Table 5**) were reported to be involved on defense response to biotic stress and other external stimuli (Ramírez et al. 2010; Depuydt and Vandepoele 2021; Gong et al. 2021). The most significant SNP was located on chromosome 3 which co-localized with Psat3g036480 that encodes for the protein kinase domain, known to be involved in the protein phosphorylation and modification amongst other cellular and metabolic processes. In general, the narrowed candidate genes have biologically related functions in response to biotic stresses, including transporter and biosynthetic processes, nucleobase-containing compound metabolic processes, transcription regulator activities, lipid metabolism, and hydrolase activity (**Supplementary Table 5**).

## DISCUSSION

Current ARR disease rating and screening methods are tedious, low throughput, time-consuming, and subjective, affecting the quality and throughput of disease scoring and subsequent screening. The conventional ARR screening in greenhouse conditions requires four to five weeks to reach the point for initiating disease evaluation, and one additional week to harvest, wash and visually rate and screen ARR. In this study, we developed a greenhouse based HTP platform that enable us to capture aerial images of field pea with the potential to capture disease symptoms quickly and objectively at symptom initiation. Using this platform, capturing images of 300 lines took 20 to 25 minutes compared to visual disease rating that required several days to complete, further highlighting the utility of HTP in a breeding program for efficient screening of thousands of genotypes. This reduction in the time needed for disease screening saves resources including greenhouse space and labor to maintain plants. Consistent with our findings, several other investigators also reported that HTP platforms equipped with cameras and sensors have successfully automated the phenotyping process and documented difficult-to-measure phenotypes quickly and objectively (Campbell et al. 2019; Mazis et al. 2020). In addition, the use of the existing greenhouse structure as a linear rail to support camera movement and image capture seemed effective and scalable. Carts could be incorporated into each necessary greenhouse room cheaply and, as the camera is removable, the camera could be taken to multiple rooms in order to collect data dependent on current project needs.

Precision, quantified through broad-sense heritability was higher for HTP-indices than visual scores. With increasing number of replications, Exg had higher heritability at 0.71-0.86 than DSI at 0.23-0.59. Thus, the visual scores require more replications (nearly two-fold) to match heritability obtained for HTP-derived indices, which is costly and requires additional time. Higher heritable variation using the HTP-derived indices can potentially increase selection accuracy. In this study, about 40% and 20% of pea genotypes were partially resistant to *Aphanomyces euteiches* based on Exg and DSI, respectively. A similar approach was taken in studies to identify and select genotypes with promising disease resistance (Oladzad et al. 2019; Zitnick-Anderson et al. 2020).

The increase in heritability can potentially increase the power of genetic analyses, improving the identification of regions associated with resistance. The number of associated SNPs for HTP-derived indices was consistently higher to the associated SNPs identified using the visual scores. GWA mapping of HTP-derived indices identified many specific loci putatively contributing to ARR resistance. All these significant loci had small effects, explaining 5 to 9% of phenotypic variance, further supporting previous studies (Ma et al. 2020; Desgroux et al. 2016) that reported the unsurprising genetic complexity of ARR resistance in pea. The underlying genes revealed fungi and biotic-stress responsive gene that are involved in catalytic activities, oxidative stress, carbohydrate metabolic processes, transporter activity, cellular processes, cell differentiation, cell growth, all of which are known to promote defense and immunity against diseases in plants. We noted important abiotic-stress responsive genes very near or directly underneath SNP associations. Psat5g280480 encodes potassium channel activity responsible for defense response and host immunity against fungi (Depuydt and Vandepoele 2021). Psat5g282800, Psat5g282880, and Psat2g167800 have been identified to play roles in biotic stimulus and defense response to oomycetes (Wakao and Benning 2005; Ramírez et al. 2010), suggesting their potential involvement in ARR disease resistance (Desgroux et al. 2017).

The SNP-level knowledge obtained through this study will help to have a fuller understanding of the causal genes conferring resistance to ARR. The identified SNP markers and genes conferring host immunity associated with the disease indices have the promise to rapidly advance cultivar development when integrated using marker-assisted selection. The identified resistant pea germplasms and genes conferring resistance could then be used to build a more durable *Aphanomyces* root rot resistance response in pea breeding programs and to help studying the genetic mechanisms associated with the disease.

One possible limitation from our study was capturing overhead canopy images from 2.5 meters above the plants using an RGB camera, which may not have given us the adequate resolution needed to trace minute ARR symptoms that appeared within the canopy area of field pea, particularly when symptoms and lesions were prevalent around the root, thus mostly under the canopy (Gaulin et al. 2007; Quillévéré-Hamard et al. 2018). Therefore, improving the phenotyping platform by reducing the focal length and building a phenotyping chamber can increase resolution and light intensity while reducing noise to improve digital indices to represent the visual scores. In addition, rapidly evolving phenotyping tools, including multispectral, hyperspectral, or thermal imaging (Marzougui et al. 2019; Sankaran, Quirós, and Miklas 2019; Hashim et al. 2020), can also be implemented along with machine learning algorithms (Shruthi et al. 2019; Mohanty et al. 2016) to further enhance the speed, precision, and accuracy of ARR phenotyping. Nevertheless, our pilot study provided valuable insights into the utility of this platform for disease evaluation and characterization both at the phenotype- and DNA-level. We are also underway testing this platform to other important diseases in pulse crops, including field, chickpea, and lentil, that show visible symptoms at the canopy level.

## CONCLUSION

In this study, we developed a greenhouse-based phenotyping platform that is objective, high-throughput, and more precise than traditional visual scoring. Using the 300 advanced breeding lines of the NDSU pulse breeding program for pilot testing this platform, precision estimated through broad-sense heritability was consistently higher for HTP-indices than visual scores, potentially increasing the power genetic mapping. GWA mapping identified a substantial number of associated SNPs for HTP-indices, which was consistently higher and with some overlap to the associated SNPs identified using the visual scores. We identified previously known genes known to be involved in the biological pathways that trigger immunity against ARR, and a few novel QTLs that may be worthy of validation in the future. The knowledge generated from this study can be further validated and used for developing a long-term, durable resistance to ARR against *Aphanomyces euteiches.*

## AUTHOR CONTRIBUTIONS

NB, JP, and PF conceived the idea and designed the study. MAB, LP, JJ, MM, JK executed the greenhouse trials. LP and RAS performed DNA extraction, constructed the library. DF, JP prepared inoculum, inoculated the trial, and visually score ARR diseases. DF, MAB captured images. JS wrote algorithms for and processed the images. MAB analyzed data, curated SNPs, and ran GWA. NB and SAA oversaw statistical analyses. MAB, HW, SAA, and NB wrote the manuscript. All authors edited, reviewed, and approved the manuscript.

## ACKNOWLEDGMENTS

The authors would like to acknowledge the funding provided by. We also acknowledge the support from USDA-NIFA (Hatch Project ND01513) and the North Dakota Department of Agriculture through the Specialty Crop Block Grant Program (19-429). Technical assistance from Didier Murillo, Ahasanul Hoque and hourly students gratefully acknowledged. This investigation used resources of the Center for Computationally Assisted Science and Technology (CCAST) at North Dakota State University, Fargo, ND, USA which were made possible in part by NSF MRI Award No. 2019077.

---

**Supplementary Figure 1.**
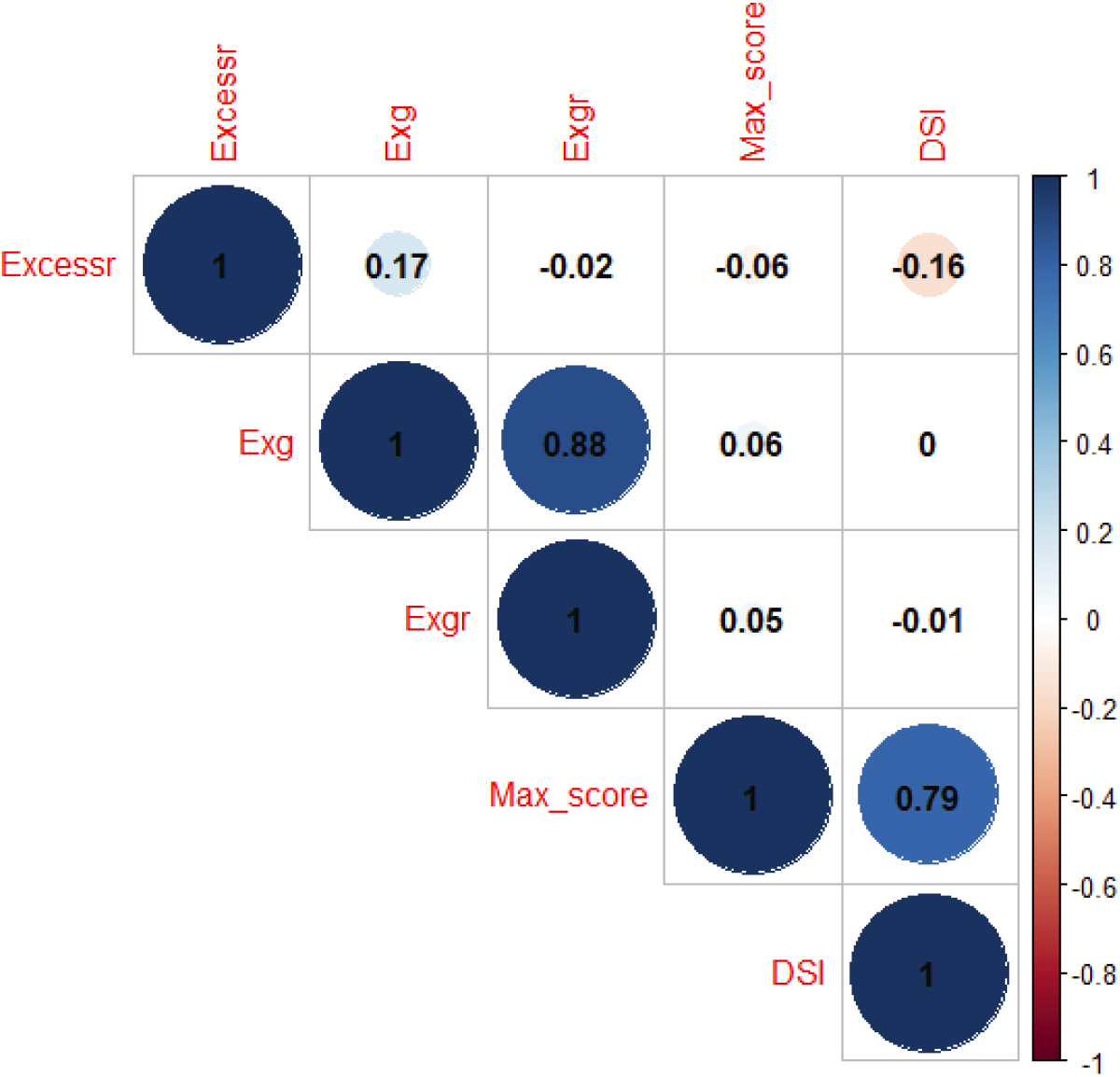
Correlation estimates of visual scores and HTP-indices across the five replications.

**Supplementary Figure 2.**
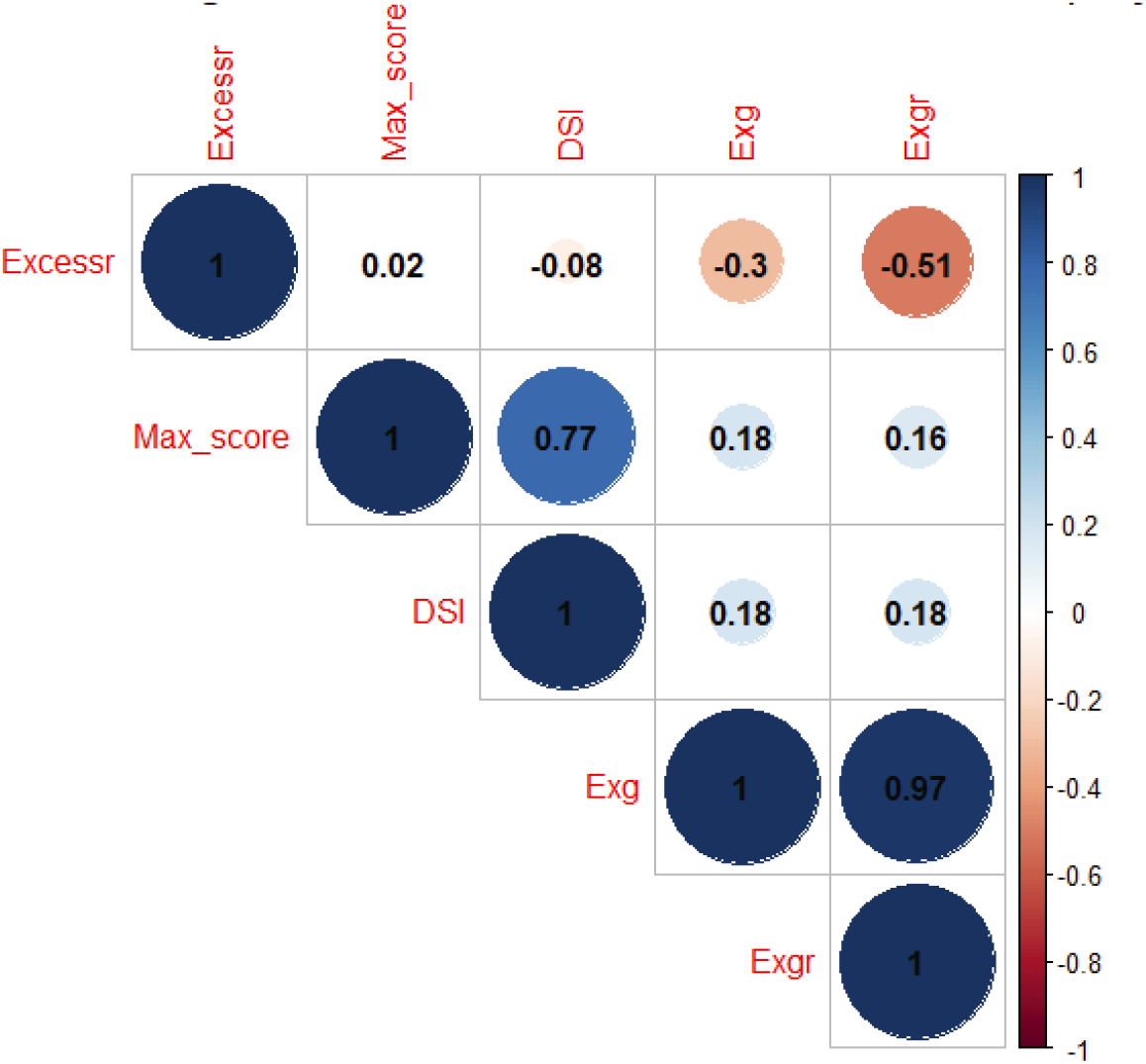
Correlation estimates of visual scores and HTP-indices in fifth replication.

**Supplementary Figure 3.**
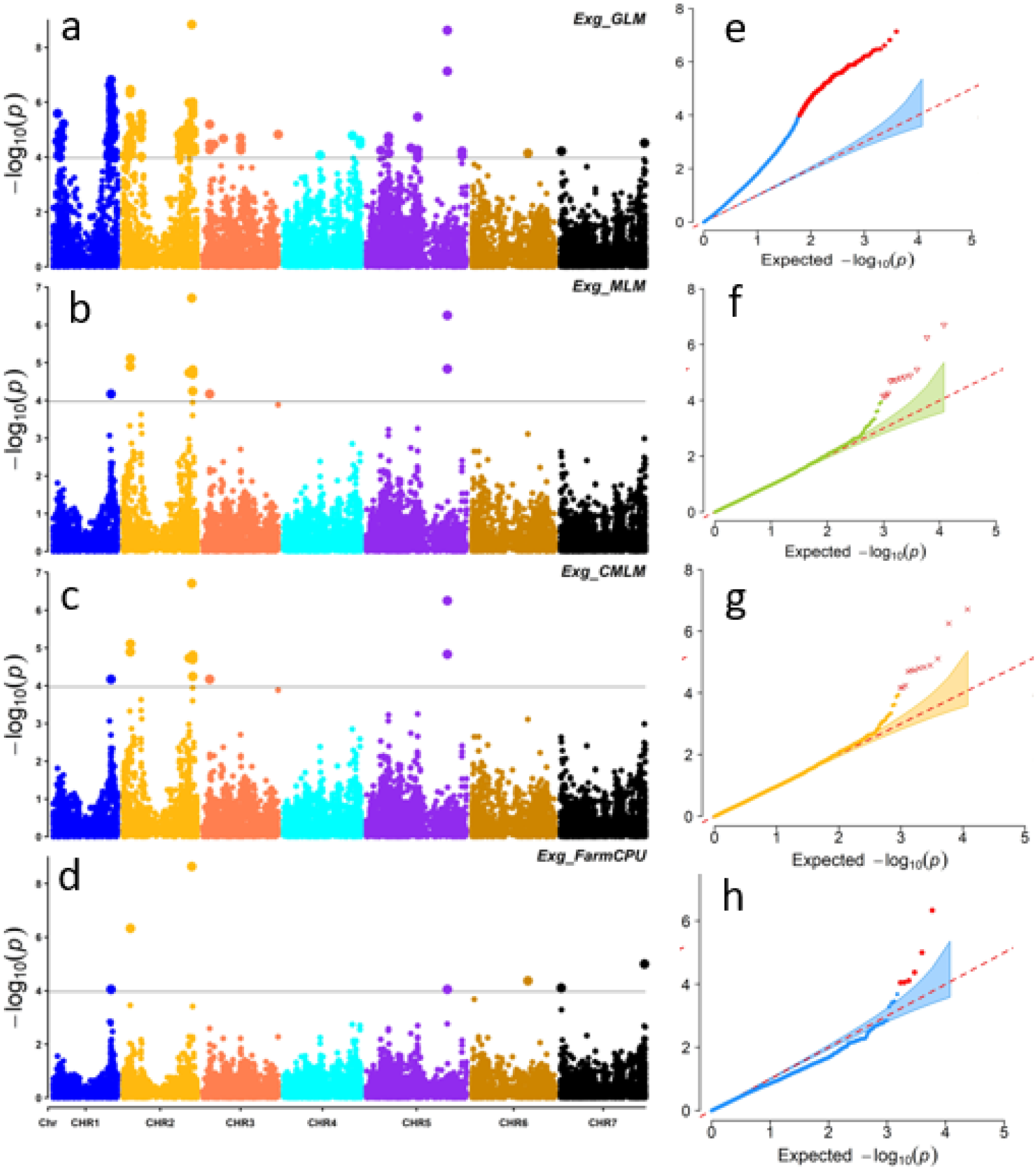
Manhattan plots for Exg exhibited associations with SNPs using a) GLM, b) MLM, c) CMLM, and d) FarmCPU that are plotted on the x-axis to their respective chromosome position on the y-axis. The horizontal line is the calculated threshold for estimating the significant association; e) Quantile Quantile-plot (QQ-plot) using GLM, f) QQ-plot using MLM, g) QQ-plot using CMLM, and h) QQ-plot using FarmCPU.

**Supplementary Figure 4.**
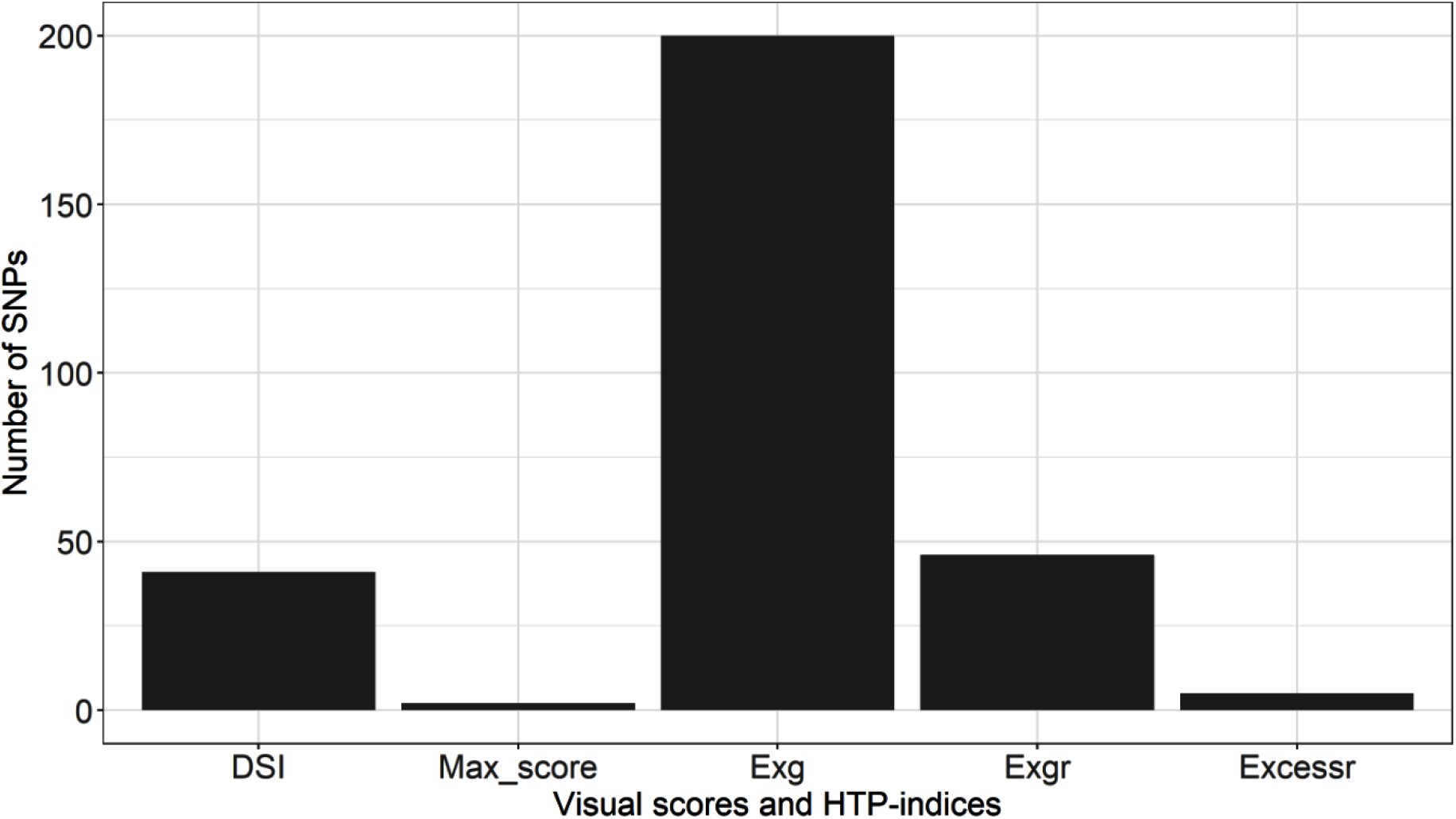
Summary of significantly associated single nucleotide polymorphism (SNPs) for visual scores and HTP-indices.

